# Whole Exome Sequencing and Single-Cell DNA Sequencing for Assessment of Clonal Heterogeneity and Evolution in Acute Myeloid Leukemia

**DOI:** 10.1101/2025.05.30.656982

**Authors:** Sadiksha Adhikari, Samuli Eldfors, Esa Pitkänen, Markus Vähä-Koskela, Caroline A. Heckman

## Abstract

**Background:** Acute myeloid leukemia (AML) progresses by the accumulation of somatic mutations and clonal expansion of pre-leukemic cells. Patients may respond to initial therapy, but often relapse, underscoring an evolving disease. Next generation sequencing technologies are being applied to AML for risk stratification and monitoring treatment response. We aimed to evaluate the efficiency of whole exome sequencing (WES) and single-cell DNA sequencing (scDNA-seq) for determining clonal heterogeneity and evolution in AML induced by treatment, assessing strengths and limitations of each technology.

**Methods:** We conducted WES and scDNA-seq on samples from 6 patients with AML, including sequential samples from four patients. We identified somatic variants, clonal composition and phylogeny using both technologies and compared the results.

**Results:** WES detected more variants and clones due to broader coverage, while scDNA-seq provided clonality results for targeted genes revealing zygosity and rare clones. Both techniques missed clinically important variants, posing challenges for clinical application. However, they identified similar founding clones and strong correlation of variant allele frequencies and clonal prevalences.

**Conclusions:** As both technologies can overlook variants, multiple technologies should be utilized to understand clonality in heterogeneous diseases such as AML. Careful scDNA-seq target panel planning, utilizing knowledge obtained from bulk sequencing, can offer more information on clonal heterogeneity.

**Graphical Abstract:** 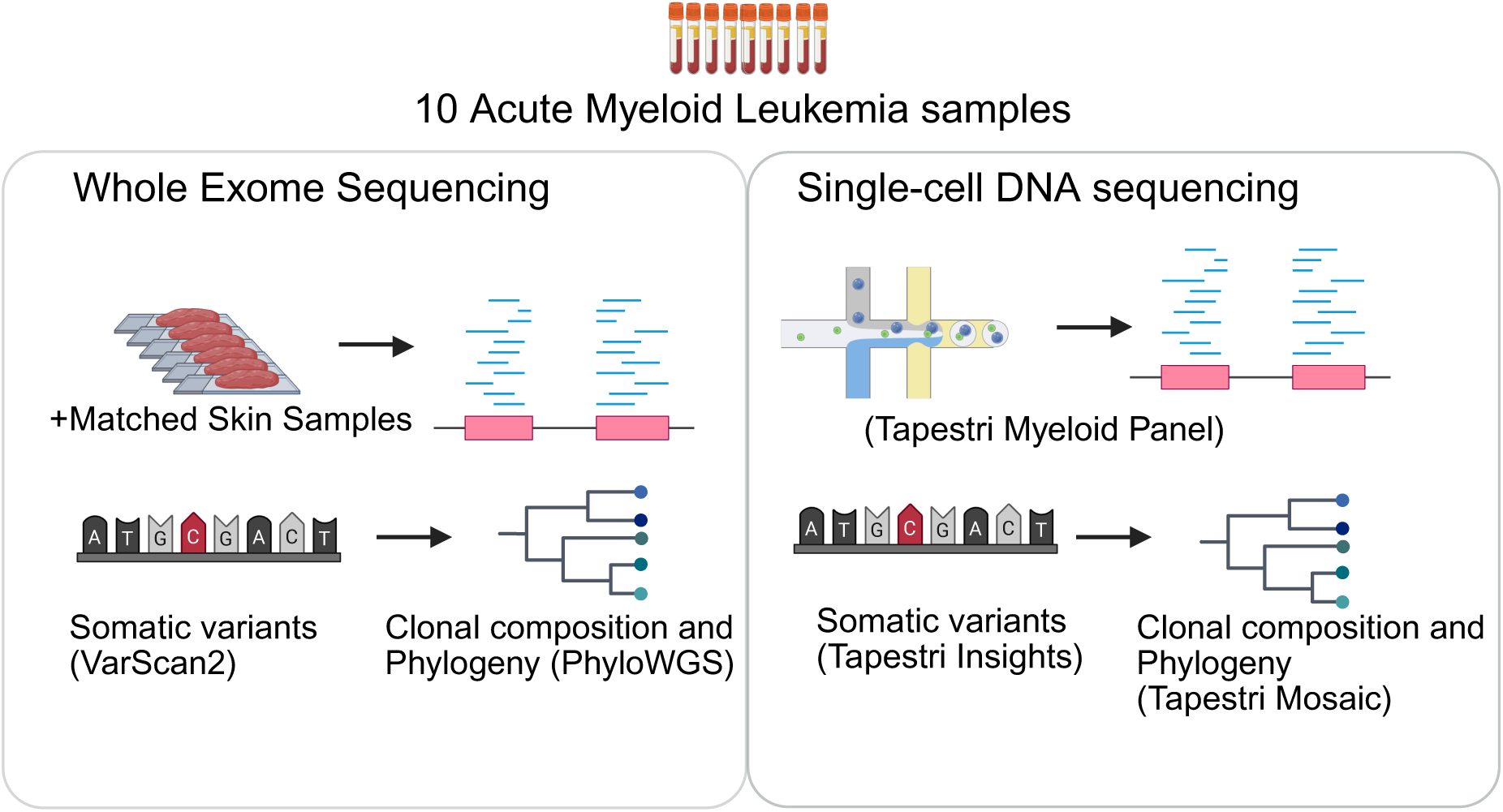

## Introduction

Acute myeloid leukemia (AML) develops by the accumulation of variants in pre-leukemic hematopoietic stem and progenitor cells, that can start occurring years before diagnosis, and result in clonal expansion [1,2]. The majority of patients diagnosed with AML initially respond to induction chemotherapy and achieve complete remission, but most will eventually relapse. This indicates an evolutionary process wherein treatment-sensitive AML cells undergo changes leading to acquired resistance, or the treatment creates a selection pressure leading to the outgrowth of a drug resistant subclone. When classified by their genetic profile, AML patients exhibit diverse variants within subtypes, allowing multiple AML subclones in a single patient [3]. Early whole genome sequencing studies have revealed two clonal evolutionary patterns of relapse: 1) the founding clone persisted after chemotherapy, but gained additional mutations, which enabled it to expand into the dominant clone at disease recurrence; or 2) a surviving subclone of the founding clone acquired additional variants and expanded at relapse [4]. Subsequent studies employing longitudinal single cell DNA sequencing and advanced subclonal cell sorting with xenografting have increased the granularity of the identification of multiple leukemic founder trajectories in individual patients and provided the basis for additional models of leukemia evolution and classification [5,6]. Clonal complexity has been linked to rapid disease development and progression [7]. While pharmacologic agents that target most recurrent AML variants are still lacking, often patients present with multiple mutations and clones, which challenges the long-term efficacy of targeted therapies. Recurrent genetic variants have been incorporated into risk stratification to predict disease prognosis and treatment responses in AML[5,8,9]. These studies underscore the importance of understanding AML as an evolutionary process with multiple clones emerging concurrently, sequentially, and adaptively in response to challenges. To this end, clinical assessment of tumor heterogeneity, rather than variants alone, can help the development of more-effective novel therapeutic strategies for addressing the disease as a whole [10].

Various methodologies are currently in use and continually emerging to identify clonality and evolution in cancer. Bulk sequencing, especially whole exome sequencing (WES), has been widely applied for accurate variant detection across gene coding regions. Bulk sequencing tools delineate subclones by grouping variants with similar allelic fractions [11,12], often adjusting the copy number at the variant loci [13,14]. The technique struggles with detecting rare variants, nevertheless, will remain crucial for clonal reconstruction due to its cost effectiveness and data availability of large study cohorts across various cancers. As an example, large, publicly available WES AML datasets were used to uncover correlations between somatic variant abundance and clonal evolution patterns with clinical patterns and drug sensitivity [7].

In contrast, single-cell DNA sequencing (scDNA-seq) has long been recognized for its potential capacity to identify rare variants and clones driving disease progression [15–18]. Recently, microfluidics-based high-throughput scDNA-seq has emerged as a potent technology for sequencing and analyzing individual cells, enabling direct observation of clonal genetic diversity and evolutionary history at a finer scale [19]. AML studies utilizing this technology have identified rare variants in relapse-prone patients [20], as well as clonal evolution, subclonal trajectories and driver variants co-occurrence, and mutual exclusivity [3,21]. Despite its potential to revolutionize clonal evolution studies, scDNA-seq currently faces technical and practical challenges, such as restrictions imposed by the chosen probe panel, allelic dropouts, amplification bias, high sequencing cost, and lack of large datasets.

The techniques are constantly evolving, with new methods emerging to provide a deeper understanding of clonal diversity and its role in various biological phenomena. Thus far, a gold standard for analyzing clonality remains elusive, and therefore, researchers must navigate available methodologies to address this analytical challenge. There needs to be more comprehensive independent assessments of these technologies to identify clonal heterogeneity and tumor phylogeny. Therefore, this study aims to assess the efficiency of intratumor heterogeneity and clonal evolution identification in AML, using WES and scDNA-seq. Additionally, the study aims to provide insights into the capabilities and limitations of each of these technologies.

## Materials and methods

### Patient material

Bone marrow aspirates and skin biopsies were obtained from patients with AML after informed consent, following protocols approved by the ethical committee of Helsinki University Hospital (permit numbers 239/13/03/00/2010, 303/13/03/01/2011) in compliance with the Declaration of Helsinki. Mononuclear cells (MNCs) were separated using Ficoll density gradient and either directly processed for sequencing or viably cryopreserved for later use. Genomic DNA was extracted from the MNCs and the skin biopsies using the DNeasy Blood & Tissue kit (Qiagen, Hilden, Germany).

### Whole exome sequencing

WES was performed on 10 AML bone marrow samples with their matched skin biopsy controls and somatic variants were called as described previously [22]. Briefly, the exomes were captured with 3 μg DNA utilizing the SureSelect Clinical Research Exome kit (Agilent Technologies, Santa Clara, CA, USA), or the SeqCap EZ MedExome kit (Roche Nimblegen, Seattle, WA, USA). The WES libraries were sequenced using the HiSeq 1500, 2000, or 2500 platform (Illumina, San Diego, CA, USA). Each sample underwent sequencing with 4×10^7^ paired-end reads of 2×50 bp read length for the germline control and 10×10^7^ paired-end reads of 2×100 bp read length for tumor samples. Reads were processed and aligned to the human reference genome GRCh37 using BWA and GATK realignment preprocessing. Somatic variants were called using Varscan2 [23]. Varscan2 assesses somatic status by comparing tumor and normal genotypes along with supporting reads using Fisher’s exact test. Variants were further filtered to include somatic calls with a Fisher’s exact *p*-value threshold of less than 0.05. The internal tandem duplication variant in the *FLT3* gene (*FLT3*-ITD) was identified using ITDetect tool [24]. Variants were recovered if misclassified as germline. Quality statistics were identified using Qualimap (v 2.3) [25].

### Clonal identification and phylogeny reconstruction by WES

Clonal identification and phylogeny reconstruction were performed using PhyloWGS [14]. PhyloWGS requires that for paired samples from same patient, the read count needs to be retrieved for each somatic variant position for all samples. Bam-readcount (version 0.8.0) [26] was used to retrieve the variant read count in sample pairs where variants were observed in one paired sample but not detected in another. Variants filtered due to *p*-value cutoffs for somatic status were retrieved with this method for accurate identification of clones and phylogeny. Allele-specific copy number and tumor purity were determined using Facets-suite (v 0.6.0) [27]. This involved utilizing tumor and normal BAM files, along with an SNP pileup file generated from common SNPs sourced from the National Center for Biotechnology Information (human_9606_b150_GRCh37p13). Somatic variant data was merged with copy number results from Facets-suite to obtain the allele-specific copy number for each variant region.

PhyloWGS was run using four MCMC chains, and other default settings. For identifying possible polyclonal seeding, --include-multiprimary --max-multiprimary 1 parameters were used. The lowest normalized log-likelihood (nLgLH) value was used to find the best phylogeny tree. Genes present on each clone were considered driver genes if they were labelled as oncogenes and tumor suppressor genes in any cancer in the OncoKB database [28], with the last update on 12/21/2023.

### Single-cell DNA sequencing

Targeted scDNA-seq data was produced with the Tapestri v2 DNA workflow (Mission Bio Support Center, 2023). The Tapestri Myeloid Panel, which targets 45 genes with 312 amplicons was applied to 10 AML samples. The sample libraries were sequenced on the Illumina NovaSeq 6000 system using read lengths 150bp (Read 1, Read2) and 8bp (i7 Index, i5 Index). Data processing and analysis - alignment, mapping, cell calling and genotype calling (GATK) – were performed through the Tapestri pipeline (DNA v 2.0.1) using BlueBee’s high performance genomics platform. Alignment was done against human genome reference GRCh37.

Downstream analysis was performed using Tapestri Insights (v2.2) and Mosaic software (v3.1.1) [29] developed and provided by Mission Bio. Variants were filtered using the following default parameters: minimum depth=10, minimum genotype quality=30, reference variant allele fraction (VAF)=5, homozygous VAF=95, heterozygous VAF=30, number of iterations = 10. Variants were required to be genotyped in at least 50% of total cells and mutated in at least 1% of the total cells. Results were manually inspected to obtain pathogenic, high-quality variants in coding regions using information from ClinVar, DANN score, and Varsome from the Tapestri insight outputs. Currently, manual inspection of variants is a required step in this analysis. Variants mutated in more than 95% of the cells were discarded as germline and if variants were detected in one Tapestri tool but not the other, such variants were provided on a whitelist, as suggested by the developer of the tools.

### Clonal identification and phylogeny reconstruction by scDNA-seq

Phylogeny was predicted using selected variants with the COMPASS tool [30], implemented within the Mosaic script, which considers cell-specific variants and copy number information to predict the phylogeny. For clonal identification, the minimum clone size was adjusted per sample from 1% to 0.1% lowering the values to identify rare clones. Clones with an allelic dropout (ADO) score above 0.8, small clones, and clones with missing information were excluded.

### Statistical calculations and visualization

Statistical calculations and visualization were performed using several packages in R. Bar plots and scatter plots were generated using the ggplot2 package. Data were filtered so that only variants and clones detected by both platforms were used for correlation analysis. Correlation analysis was performed with the ggpubr package using the Pearson correlation coefficient. The Shannon index, indicative of clonal dominance, was calculated from clone proportions using vegan package (v 2.6-4). Fishplots were created using fishplot package (v 0.5.2). Missed variants were investigated using IGV (v 2.16.0) [31]. GenVisR (v 1.34.0) was used for WES genome-wide copy number variation (CNV) plot. Phylogeny plots were created with BioRender (BioRender.com).

## Results

We evaluated the performance of WES and scDNA-seq to identify somatic variants and clones, and to construct phylogenies. WES was performed on bone marrow aspirates with matched skin biopsies for six AML patients. ScDNA-seq was performed using bone marrow aspirates from the same samples. The workflow is shown in **Figure 1**. The main baseline characteristics of the study population are summarized in **Table 1**.

**Figure 1.**
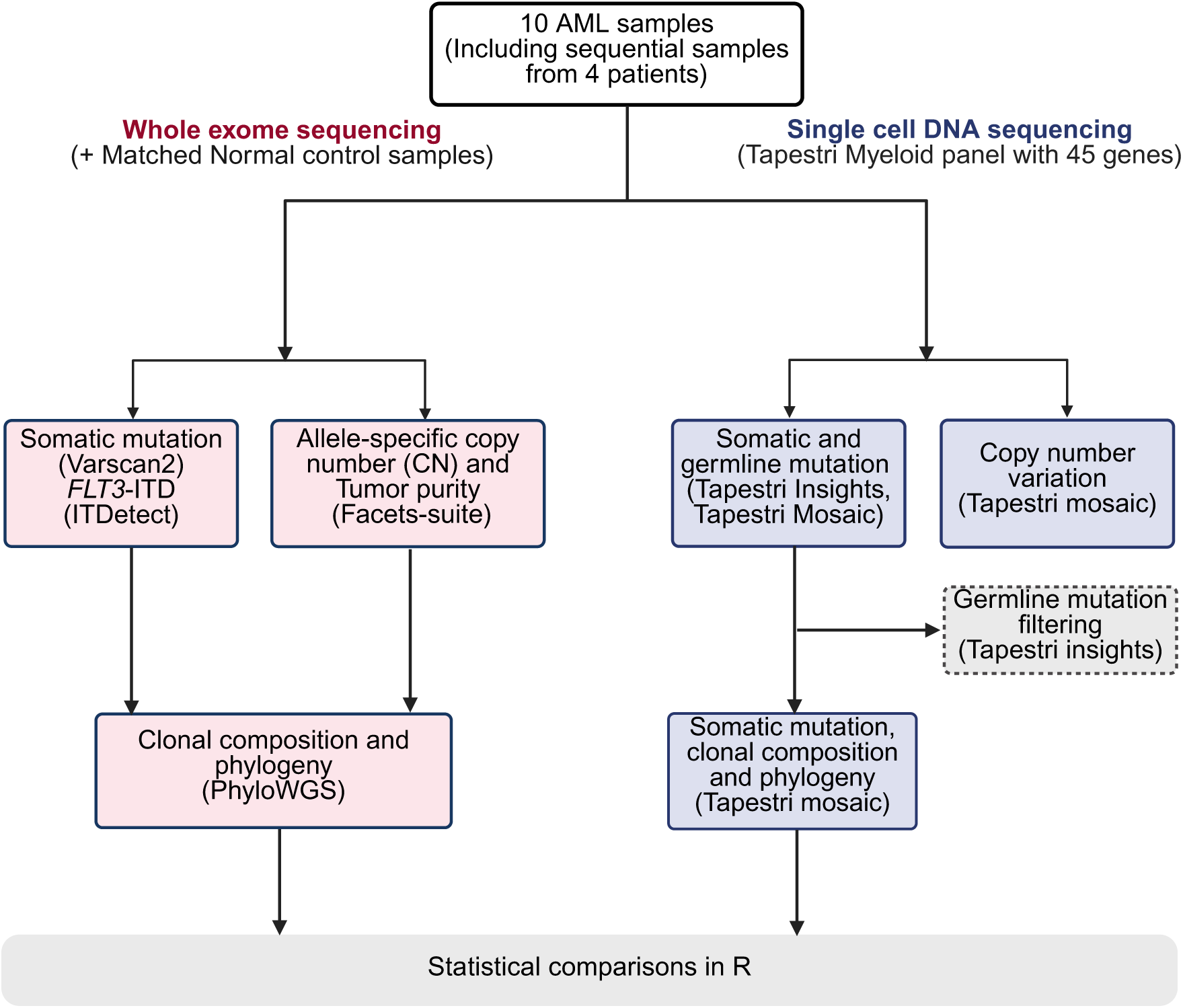
Workflow of the project.

**Table 1.** Clinical characteristics of the AML patient cohort.

| Sample | Sex | Age, y | Cytogenetics | ELN 2022 classification | Disease status | FAB Subtype | Treatment received | ASCT |
| --- | --- | --- | --- | --- | --- | --- | --- | --- |
| FPM_AML_112_01 | Male | 64 | Normal, ASXL1 mut | Adverse | Diagnosis | FAB M1 | No (Treated for thromb) | No |
| FPM_AML_112_02 | Male | 64 | Normal, ASXL1 mut |  | Relapse | FAB M1 | Yes | No |
| FPM_AML_131_02 | Female | 71 | Not known |  | Relapse | FAB M5 | Yes | No |
| FPM_AML_162_01 | Male | 70 | del(5q) | Adverse | Diagnosis | FAB M2 | No (Treated for essent) | No |
| FPM_AML_162_02 | Male | 71 | del(5q) |  | Refractory | FAB M2 | Yes | No |
| FPM_AML_015_05 | Male | 56 | t(1;5;14), FLT3-ITD mut | Intermediate | Diagnosis | FAB M5 | No | No |
| FPM_AML_015_02 | Male | 57 | t(1;5;14) |  | Refractory | FAB M5 | Yes | Yes |
| FPM_AML_035_01 | Female | 22 | Complex karyotype (-2, abn(5), -6q, -7, -11, -12, +13, -17, -18, +21, -22) | Adverse | Diagnosis | FAB M1 | No | No |
| FPM_AML_078_01 | Male | 69 | Not known | Adverse | Diagnosis | FAB M4 | No | No |
| FPM_AML_078_02 | Male | 70 | Not known |  | Refractory | FAB M4 | Yes | No |

### Overview of data by WES and scDNA-seq on AML samples

In general, for WES there was an average of 126,417,400 reads per sample. On average 98.3% reads mapped to genome. The mean on-target ratio was 67.9 (range 39.1%-81.9%), which is typical for this technology (**Supplementary Table 1**).

With scDNA-seq, a total of 73,490 cells and 7,349 cells per sample on average were sequenced. The mean number of reads per cell was 27,511. The mean on target ratio was 88.3% (range 81.6-90.7). The mean ADO rate was observed to be 12.6%, varying between 8% and 19.2%. Following filtering for ADOs, false positives, and missing variants, 16,307 (22%) cells out of total cells, were available for analysis to identify clonal composition; on average 1,705.4 cells were analysed per sample. Comprehensive details are presented in **Supplementary Table 1**.

### Genetic alterations identified in AML samples via WES and scDNA-seq

WES revealed 692 somatic variants, including single nucleotide variants, and short insertion and deletions (indels) in 461 unique genes (**Figure 2a**). For scRNA-seq, a total of 1,024 variant events were identified through the Tapestri pipeline. Upon filtering for false positives and germline variations, scDNA-seq identified 33 events, entailing 16 unique somatic variants across 12 unique genes. The large difference in variant capture was due to the significantly larger library size of capture targets of WES (over 41 Mb) compared to the scDNA-seq myeloid gene panel (1.5kb). Therefore, for direct comparison between variants detected by both WES and scDNA-seq, only the forty-five genes exclusively encompassed within the Tapestri targeted panel were considered (**Supplementary Table 2**). Following filtration, WES detected 33 events, comprising 19 unique variants across 14 unique genes. Notably, two cases, namely sample FPM_AML_131_02 and FPM_AML_035_01, exhibited an absence of variants in the myeloid panel genes (**Figure 2b**). In total, variants across 14 genes were discerned in eight samples. 27 events with 15 unique variants were found to be shared between the datasets (**Figure 2c**).

**Figure 2.**
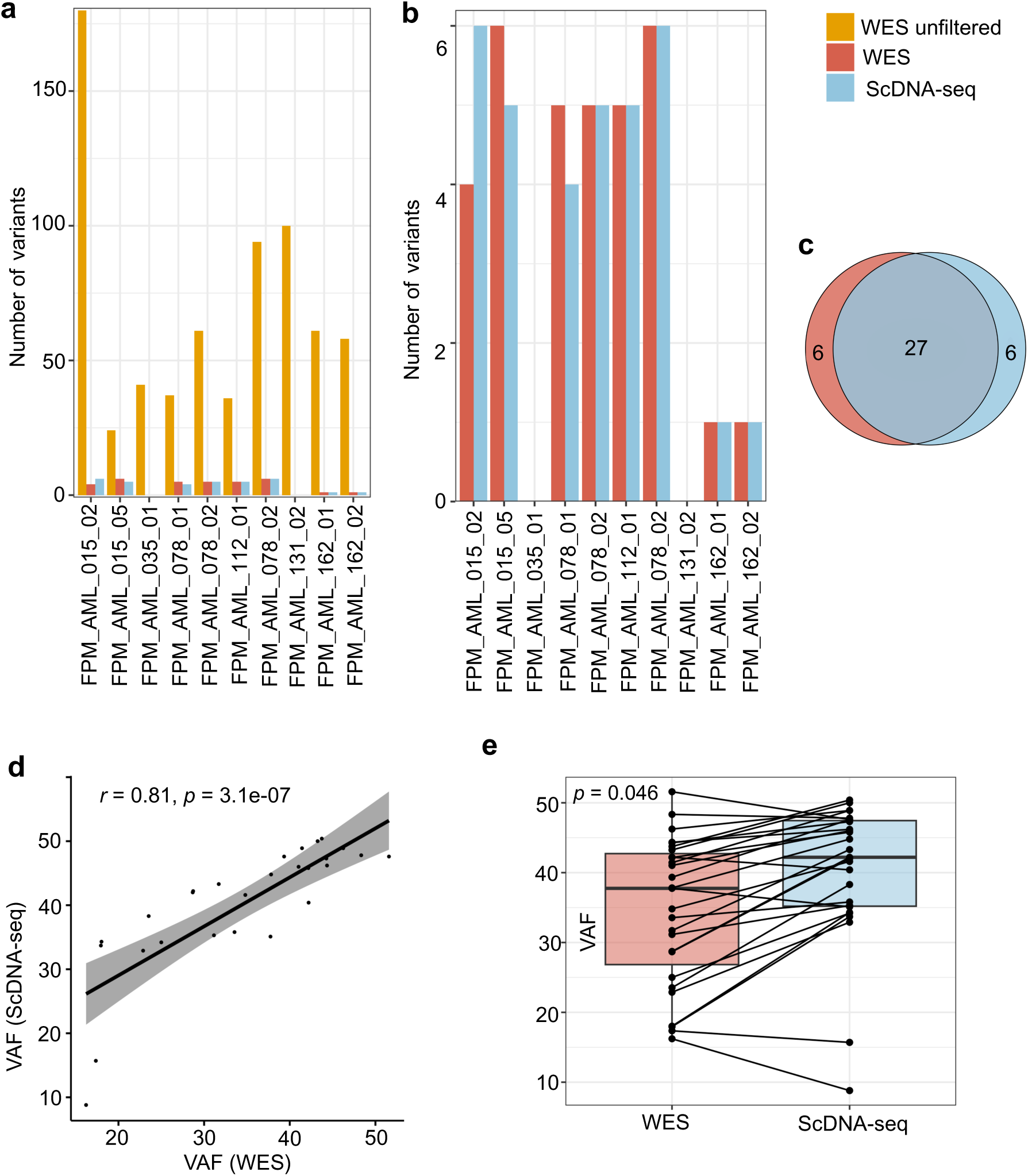
Variants identified by WES and scDNA-seq across 10 AML samples. (**a**) The total number of somatic variants detected by WES, filtered WES variants (only genes included in Tapestri Myeloid panel), and total somatic variants detected by scDNA-seq. (**b**) Total variants in genes included in the Myeloid panel for WES and scDNA-seq (**c**) Venn diagram showing common events identified in WES and scDNA-seq. (**d**) Correlation between variant allele frequency of common variants detected by WES on the *x*-axis and scDNA-seq on the *y*-axis. (**e**) Box plot comparing variant allele frequency of common variants detected by WES and scDNA-seq using paired Wilcoxon-test.

Variants such as *ETV6, PHF6*, and *IDH2* were occasionally overlooked by WES but were identified by scDNA-seq. Further analysis revealed that the WES pipeline excluded *ETV6* because its *p*-value, determined by Varscan2’s Fisher’s exact test for somatic status, was marginally above the cutoff of 0.05 (*p*=0.053). In contrast, variants in *PHF6* and *IDH2* were not detected because they were presented with low VAF. Although *NF1* and *BRAF* were included in the Tapestri Myeloid Panel, those variants’ positions were not covered by the scDNA-seq panel’s amplicons, hence, were not reported by scDNA-seq. In contrast, despite the variant position being included in the panel, scDNA-seq failed to detect mutation to *ASXL1*. When the *ASXL1* variant position was provided as a whitelist based on the variant position from WES, the results confirmed the *ASXL1* variant present in a clone with *DNMT3A*/*IDH2*/*JAK2* mutations, mirroring WES results. However, we observed that over 81% of the cells had insufficient genotyping data in the variant position and was therefore filtered out by the Tapestri analysis software (**Supplementary Table 3**). Despite the discrepancies, the VAF of common variants exhibited a strong and significant positive correlation (r=0.81, *p*=3.1e-07) (**Figure 2d**). ScDNA-seq was slightly more sensitive in obtaining VAF compared to WES (*p*=0.046, paired Wilcoxon-test) (**Figure 2e**).

### Clonal prevalence and diversity identified by WES and scDNA-seq

Comparing the number of clones, WES yielded more clones per sample (median 6.5) than scDNA-seq (median 2.5) shown in **Table 2**. This was expected, as WES covers a substantially larger spectrum of variants. Next, the cellular prevalence of clones between the datasets was compared, detailed in **Supplementary Table 4**. Six out of eight samples exhibited dominant clones with co-occurrent variants among the common variants identified by WES and scDNA-seq, implying consistency in identifying major clonal populations. However, variability in cellular prevalence percentages was observed, possibly influenced by the wildtype population. Notably, the Tapestri tools determine the wildtype proportion directly from cell numbers, whereas PhyloWGS infers wildtype content from the cancer cell fraction of somatic variants. Despite this methodological difference, a positive correlation between cellular prevalence in the datasets (r=0.91, *p*=7.3e-06) was observed (**Figure 3a**). Furthermore, the Shannon index, calculated from clone proportions, was compared across datasets. This index provides insight into clonal diversity by considering both the number of clones present and their relative abundances. By paired Wilcoxon test on only the clones present in both datasets, WES exhibited significantly higher diversity within the AML cohort compared to scDNA-seq within the same dataset *(p*=0.0078) (**Figure 2b**).

**Figure 3.**
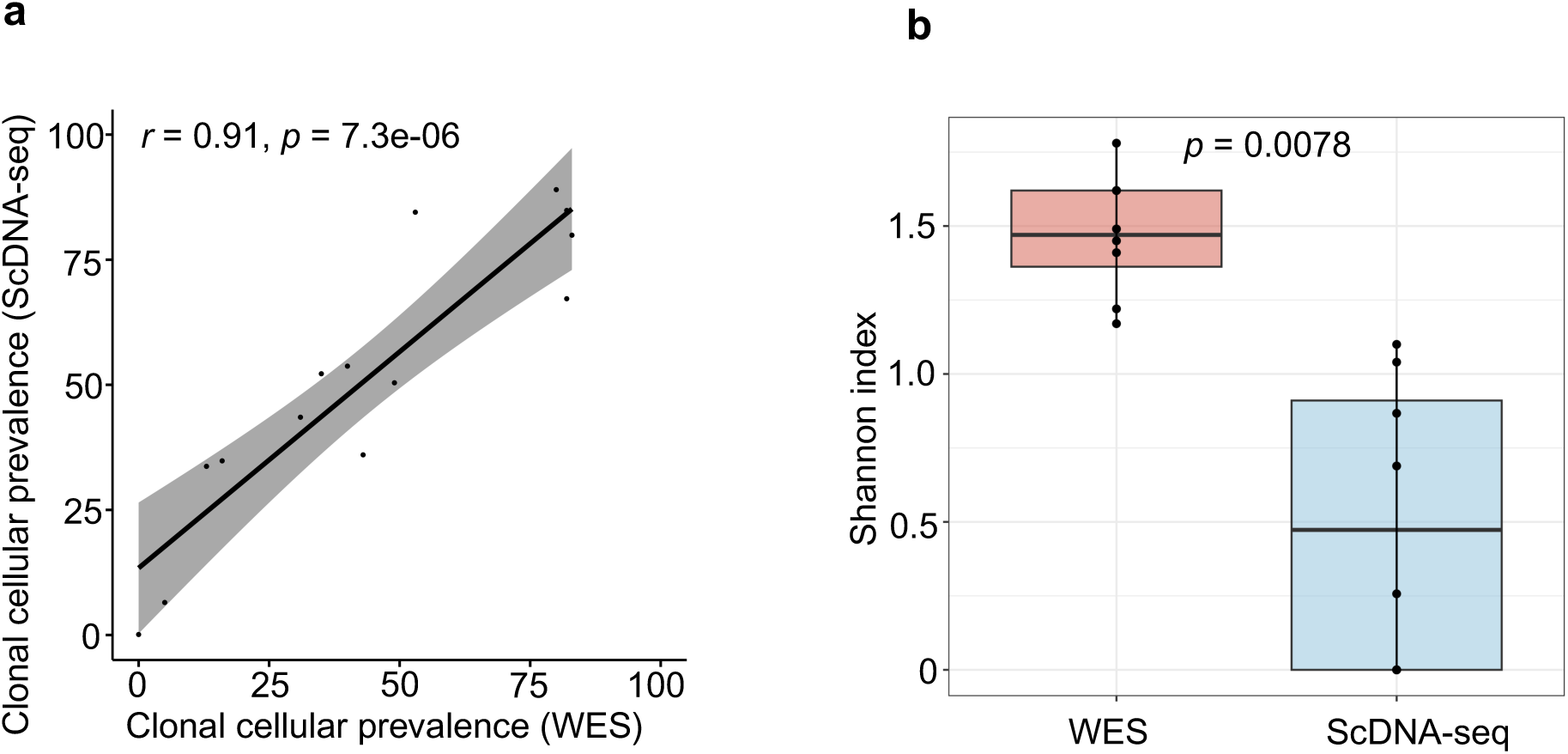
Clonal prevalence and diversity. (**a**) Correlation of cellular prevalence between clones common in each sample between WES and scDNA-seq. Total clonal prevalance was calculated by combining subclones. (**b**) Box plot comparing Shannon diversity index calculated for each sample using cellular prevalence values of WES and scDNA-seq. For paired Wilcoxon test, only clones present in both datasets and only paired samples were considered.

**Table 2.**
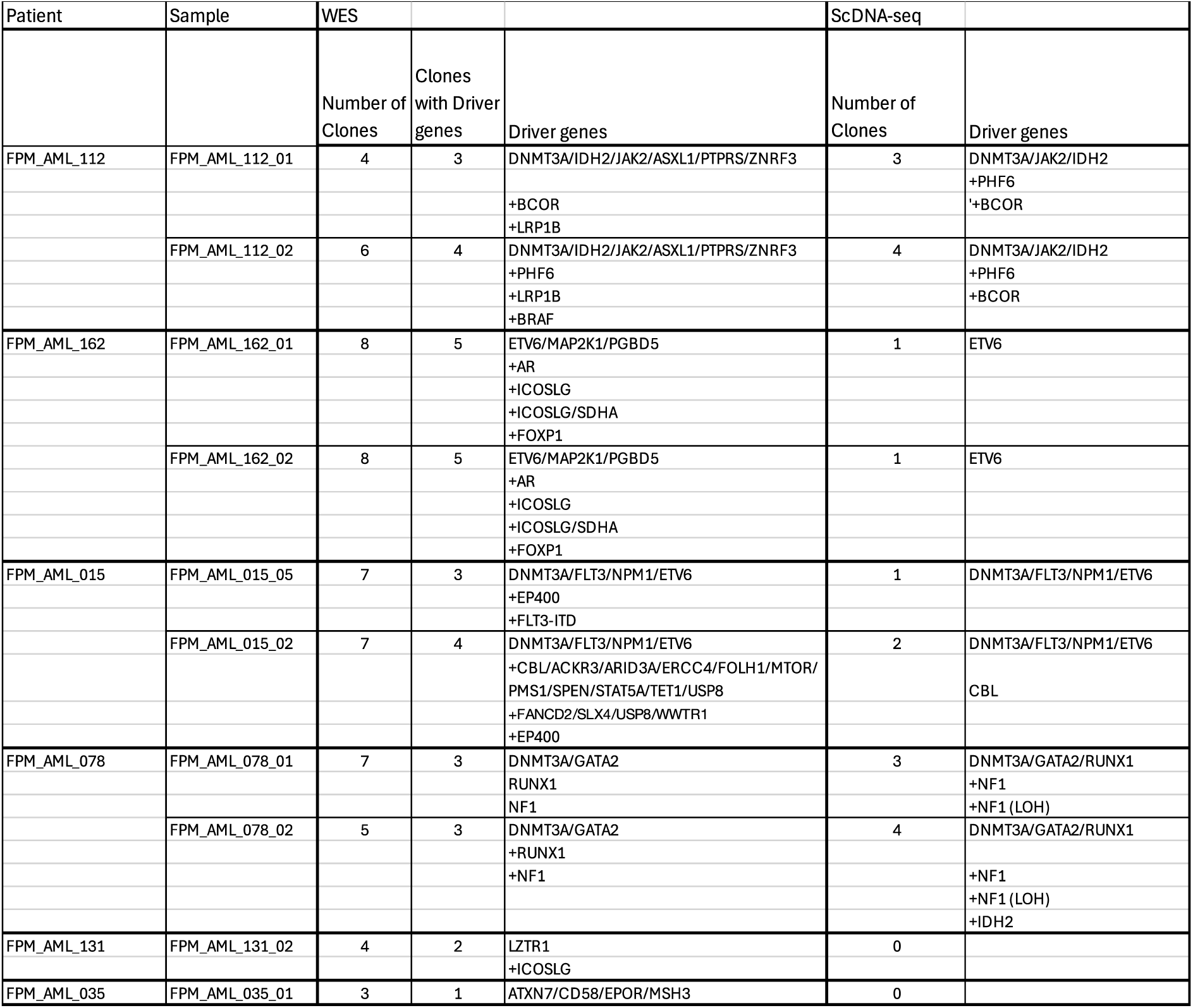
Summary of clonal numbers, populations and associated driver genes identified through WES and scDNA-seq in each sample.

### Complementary insights into clonal composition and trajectories by WES and scDNA-seq

Diverse clonal evolution patterns were evident from both WES and scDNA-seq in individual cases (**Figure 4**). In the case of patient FPM_AML_112, both WES and scDNA-seq identified the *DNMT3A*/*IDH2*/*JAK2* mutant clone as dominant (**Figure 4a, b**). ScDNA-seq identified linear evolution of clonal events as well as branches. A rare *PHF6 (R274Q)* subclone with only 9 cells at diagnosis gained an additional subclone with another variant of *PHF6 (K241Nfs*24*) and expanded to almost 40% of the population at relapse. Additionally, scDNA-seq also revealed a *BCOR* clone at diagnosis, which collapsed to only 2 cells at relapse, showcasing the unique contribution of scDNA-seq for deciphering phylogeny and clonal evolution over time. It is important to note that these rare clones present in few cells were not observed when the samples were analyzed individually (data not shown). WES identified all this as branching evolution and missed the *PHF6 (K241Nfs*24*) subclone. In contrast, WES revealed a clone bearing a *BRAF* mutation co-existing with the *PHF6* clone, which scDNA-seq failed to detect because of insufficient coverage of the *BRAF* gene. ScDNA-seq identified homozygous deletion in *RUNX1* in all mutant clones **(Supplementary Figure 1a-b**). RUNX1 is present in chromosome 21; WES identified loss in chromosome 21 in relapse sample **(Supplementary Figure 2**) but was unable to detect this loss in any clones. This could be because bulk tools utilize allele-specific copy number only for regions with somatic variants to identify clones. If there is no variant in a mutant region, precise copy number changes will be challenging to be reported at clonal level.

**Figure 4.**
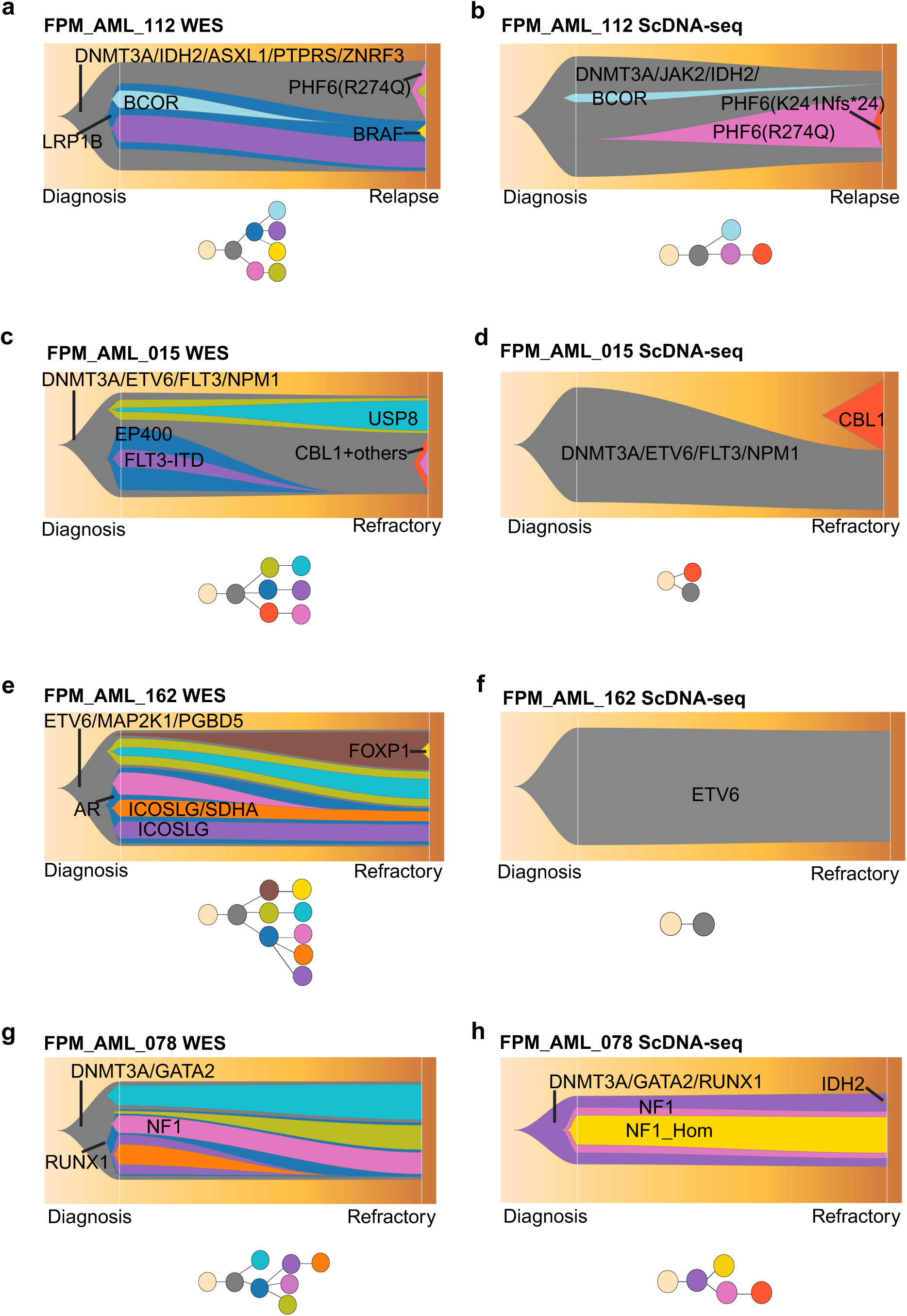
Clonal prevalence over time of four patients identified with WES (left) and scDNA-seq (right). Predicted phylogeny for each patient are provided below the fishplot. Patient-wise, the same clones identified with WES and scDNA-seq are indicated with the same color.

Patient FPM_AML_015 exhibited notable differences in clonal evolution results. ScDNA-seq indicated a clonal switch from the *DNMT3A/NPM1/FLT3/ETV6* mutant clone at diagnosis to a coexisting *CBL* mutant clone (**Figure 4d**). However, WES revealed *DNMT3A/NPM1/FLT3/ETV6* as a founding clone with subclonal branches, one being the *CBL* clone along with numerous driver genes and over a hundred other variants, dominating half of the clonal population in the refractory sample (**Figure 4c, Supplementary Table 4**). Interestingly, both WES and scDNA-seq identified a gain in chromosome 8 in the refractory sample only in the *CBL* clone (**Supplementary Figure 2, Supplementary Figure 1d**). The patient had undergone allogeneic transplantation, suggesting that the clone with *CBL* and other genes might have originated from the donor rather than indicative of patient-specific clonal evolution. WES identified a low-frequency subclone with *FLT3-*ITD at diagnosis. ScDNA-seq failed to detect the *FLT3*-ITD clone, even when the position of the variant was whitelisted. The ITDetect tool that was used for *FLT3*-ITD detection in WES data, was also applied to scDNA-seq data. However, the process was not completed within a 48-hour period and was subsequently terminated.

In case FPM_AML_162, scDNA-seq detected only an *ETV6* mutant clone attributed to a single variant from the Tapestri panel genes (**Figure 4f**), with a copy number profile similar to the *ETV6* wildtype clone **(Supplementary Figure 1e-f**). WES identified additional subclones, with the dominant clone characterized by variants in *ETV6* with other genes known as drivers in other cancers, but not in AML, *MAP2K1* and *PGBD5* (**Figure 4e**). WES identified chromosome 5 deletion, which was also reported in the clinical karyotyping data, however, did not place it in any clones. Other copy number changes identified by WES included gains in chromosomes 3, 15, 16, 19, 20, and 22 within the dominant clone.

For patient FPM_AML_078, WES identified *DNMT3A/GATA2* as the founding clone, later acquiring subclonal variants in *RUNX1* followed by *NF1* (**Figure 4g**). In contrast, scDNA-seq revealed dominance of a clone with mutations to *DNMT3A*, *GATA2*, and *RUNX1*, followed by subclonal variants in *NF1* (**Figure 4h**). Furthermore, scDNA-seq detected a rare clone with an *IDH2* variant at the refractory stage, which WES failed to detect. Due to low prevalence (1.26%), this clone is unlikely to be responsible for disease progression, however, it was already present in 5 cells out of 6417 genotyped at diagnosis. Additionally, scDNA-seq uncovered a subclone with a homozygous *NF1* variant, corroborated by CNV analysis indicating copy number deletion of *NF1* (**Supplementary Figure 1g-h**). ScDNA-seq also identified homozygous deletion of *BCOR* in all mutant clones. At the clonal level, WES missed this CNV as *BCOR* lies in chromosome X, and the CNV-tool, facets-suite, did not report copy number variations in chromosomes X and Y.

For two samples, FPM_AML_131_02 and FPM_AML_135_01 scDNA-seq did not provide clonality information due to absence of variants in targeted genes and WES identified clones with genes that are not known as drivers in AML. Sample FPM_AML_035_01 was from a young patient, with complex karyotype, indicating that AML drivers, in these cases, are likely to be factors other than somatic variants, such as somatic structural changes. WES was able to identify copy number changes in this patient (−2, −6q, −7, −11, −12, +13, −17, −18, +21), except deletion of chromosome 22 (**Supplementary Figure 2**).

## Discussion

Our study aimed to assess the efficacy of two prominent technologies, WES and scDNA-seq, in analyzing clonal reconstruction and evolutionary trajectories in AML samples. Clonal heterogeneity analysis typically involves variant identification, followed by deciphering clonal composition and identification of driver genes in the clones. In our results WES detected a higher number of variants due to its significantly larger library size compared to the 45 myeloid gene panel used for scDNA-seq. Despite this difference, both technologies were capable and had individual strengths in deciphering clonal composition and evolution. ScDNA-seq presented remarkable precision in detecting low frequency variants, in as few as two cells, offering deeper insights especially in the clonal dynamics of longitudinal samples. Notably, these exceedingly rare clones were overlooked when the samples were analyzed individually. On the other hand, WES outperformed scDNA-seq in capturing clonal diversity, attributed to its broader genomic coverage, presenting a higher number of identified clones; this is significant as the clonal diversity index accounts for both number and abundance of clones. These findings highlight the distinct strength of the technologies and the significance of longitudinal samples for precise and consistent tracking variants over time and within individual clones. However, both technologies presented challenges with variant detection, leading to incomplete assessments of clonal composition.

Inherent limitations of WES include difficulty in capturing clones with rare variants, which can significantly impact clonality studies. For example, not identifying small clones until relapse can falsely indicate branching rather than linear evolution. Furthermore, numerous cancer studies over the past decade have shown that resistant clones typically stem from acquired resistance, i.e., the selective expansion of pre-existing variants during treatment [32,33]. Therefore, WES can overlook low-frequency variants, which often drive heterogeneity and resistance to targeted therapies, losing vital data crucial for clinical applications. A small number of AML leukemic cells that remain in the patient even after treatment, when the patient is in complete remission, referred to as minimal residual disease (MRD), is an important prognostic factor in AML because its presence is associated with a higher risk of relapse and requires techniques that can detect leukemic cells at low concentrations. Due to the capacity to detect such rare clones, scDNA-seq, combined with immunophenotyping, has been recently shown to enhance MRD detection [18,34]. Another challenge with clonal heterogeneity identification with bulk sequencing is the selection of the most effective computational tool. Inconsistencies in results from different tools have been an issue for a long time [35–37]. We also initially opted for two widely used methods, PyClone [13] and PhyloWGS [14]. However, PyClone provided only clones, did not provide phylogeny and failed to put many of the mutations into clones; we were unable to get phylogeny results with additional R packages either. Bulk method of clonality detection is labor intensive with inclusion of multiple tools and manual inspection, posing challenges for application in large cohorts.

Challenges in scDNA-seq arise due to lack of prior knowledge about variants in samples, ADOs, and incomplete gene coverage. A large proportion of cells were discarded in the analysis before the identifying phylogeny as they lacked information in one or more of the selected variants. One reason for that is ADOs, occurring due to failure to amplify one or both heterozygous alleles, which can lead to incorrect inference of genotyping [38]. When samples lacked variants in the target panel genes, no additional information was extractable regarding clones or ploidy. Additionally, incomplete gene coverage can pose challenges in clinical applications. Sequencing whole exonic reads with scDNA-seq is technically possible but prohibitively expensive for research labs, and the issue of ADO magnifies with large single-cell data. ScDNA-seq offers the flexibility to recover variants filtered out by default pipeline and tool settings by utilizing a whitelist of variants. Depending on the research or clinical goals and research settings, if variants in a cohort are identified using bulk methodologies first and then the existing scDNA-seq disease-specific panels can be enhanced by incorporating corresponding amplicons. This augmentation can potentially yield more precise data on clonal phylogeny and reconstruction, especially regarding driver variants. Additionally, utilizing the whitelist feature of scDNA-seq can be advantageous, enabling assessment of clonal composition and cell-level information based on clinically relevant genes identified through bulk sequencing. Improvements in single cell technologies, plus reducing the ADO rate and costs, can offer wider applications to larger cohorts and more clinical utility of the technology.

ScDNA-seq overlooked clinically relevant mutations to *ASXL1* and *FLT3*. Despite the high resolution of the technology, an *ASXL1* variant was genotyped in only 18% of cells, and therefore was filtered out by the pipeline. Interestingly, such low genotyping of this gene is consistent with results on this platform shown by previous studies [18,20] (Ediriwickrema et al., 2020; Guess et al 2022). The presence of an *ASXL1* variant places patients into an adverse risk group according to the ELN 2022 guidelines [8]. Thus, losing this information is unhelpful in a clinical setting. Similarly, *FLT3*-ITD variants places patients into an intermediate risk group as it is associated with higher relapse rates, reduced overall survival, and significantly impacts disease management [8,9] .Patients with a *FLT3*-ITD often require aggressive therapies and benefit from targeted *FLT3* inhibitors [39]. While next generation sequencing-based detection of *FLT3*-ITD is challenging and typically demands additional software, the scDNA-seq Tapestri pipeline integrates *FLT3*-ITD detection steps, showing past success in uncovering this variant [40]. Our results show that the challenge of *FLT3*-ITD detection by next generation sequencing persists, even with high resolution sequencing of single cells.

Another crucial aspect of clonal composition is the identification of CNVs, where scDNA-seq exhibited notable superiority. This technology not only provided copy number details, but also distinguished zygosity, identifying clones with copy-neutral loss of heterozygosity and homozygous deletions. Unlike previous DNA amplification methods, scDNA-seq was able to integrate variant and copy number alteration data into phylogenetic algorithms from single-cell data, marking a substantial progression in this field [30]. Nevertheless, it is pertinent to note that the clonal CNV data is incomplete, being limited to the targeted regions. Although WES provides genome-wide CNV data, only a few bulk processing tools like PhyloWGS offer combined subclonal copy number information and variant details. Additionally, input clonal CNV data remains partial, as clonality tools traditionally utilize only regions with somatic variants. In our dataset, loss-of-heterozygosity events detected at the genome-wide level by WES were not evident in specific clones due to exclusion of non-mutated CNV regions from the input files, or non-reporting by CNV tools when CNV was located on sex chromosomes. Nonetheless, there was a concurrence in the assignment of diploid stage and other copy numbers between technologies; for instance, a chromosome 8 deletion was consistently observed across the same clones using both scDNA-seq and WES technologies.

One limitation of this study was the potential acquisition of biases stemming from differences in library preparation and analysis pipelines, which were not accounted for in our results. Computational challenges are magnified when comparing different technologies such as WES and scDNA-seq due to lack of a standardized pipeline applicable to both methods. Nevertheless, our study assessed their respective capabilities in analyzing clonal heterogeneity while acknowledging inherent differences in library preparation and analysis pipelines. Despite some missed variants, the strong correlation of common variant VAFs and cellular prevalences between the datasets suggests consistent and reliable results between the two methods. The small sample size may impact the statistical power and generalizability of the findings, warranting a cautious interpretation. Additionally, technical constraints include the limited number of methodologies explored and the absence of comparative alternative scDNA-seq approaches. Furthermore, the research was limited to just two timepoints for longitudinal samples. More comprehensive sampling throughout a patient’s progression might capture more accurately the clonal dynamics potentially overlooked in our analysis. Lastly, while the WES coverage in this study is typical for detecting variants in AML, higher coverage might provide deeper insights into clonality. Further studies with larger patient cohorts, incorporating both technologies, could address the gap in our knowledge regarding genetic heterogeneity of AML samples, clonal evolution in response to chemotherapy, and possibly identifying common features for targeted therapies and clinical applications.

In conclusion, our study shows that the selection between WES and scDNA-seq hinges on the specific requirements of clonal analysis in the disease studies. WES is advantageous for extensive clonal analysis across large datasets, identifying driver events and their clinical correlations. However, its limitation in detecting variants with low VAF makes scDNA-seq preferable for exploring rare clones and disease progression in conditions with well-characterized driver variants. ScDNA-seq can also provide clonal copy number events such as LOH in the selected genes. Despite its challenges in genotyping certain variants and ADOs, the effectiveness of scDNA-seq can be enhanced through strategic target panel planning based on knowledge from bulk sequencing data. Such an approach can be beneficial in dissecting clonal heterogeneity and phylogeny in a heterogeneous disease like AML. Factors such as cost and the need for clonal protein-level data, which were not considered here, also play critical roles in choosing the appropriate technology for specific research goals and context.

## Acknowledgements

We are extremely thankful to the patients who donated samples for the study and the Finnish Hematology Registry and Biobank (FHRB) for providing the samples. We are thankful to Minna Suvela for technical support and patient sample processing. The authors acknowledge the assistance and support of the personnel at the FIMM Sequencing and Single-Cell Analytics units particularly for their help with library preparation and sequencing. FIMM IT services are acknowledged for providing cluster nodes to run the analysis. We also thank personnel at Mission Bio for providing data analysis training to run Tapestri tools and providing advice.

## Author contributions

S.A., E.P., C.A.H., and M.V.K. contributed to the conception or design of the study. All authors contributed to data acquisition, analysis, or interpretation; C.A.H., and M.V.K contributed to data acquisition, S.E. developed the WES pipeline, S.A. contributed to data analysis and writing. C.A.H., E.P., and M.V.K. supervised the project and helped write the manuscript. All authors reviewed the drafts and approved the final version. All authors agreed to be accountable for all aspects of the work in ensuring that questions related to the accuracy or integrity of any part of the work are appropriately investigated and resolved. The corresponding author had full access to the study data and accepts final responsibility for the decision to submit for publication.

## Ethics approval and consent to participate

The protocols used in this study were approved by an ethical committee of Helsinki University Hospital (permit numbers 239/13/03/00/2010, 303/13/03/01/2011). The study was performed in accordance with the Declaration of Helsinki. Samples were obtained from patients after receiving written informed consent.

## Consent for publication

Not applicable.

## Data availability

The datasets generated during and/or analyzed during the current study are not publicly available due to privacy and ethical limitations but are available from the corresponding author on reasonable request.

## Supplementary figures

**Supplementary Figure 1.**
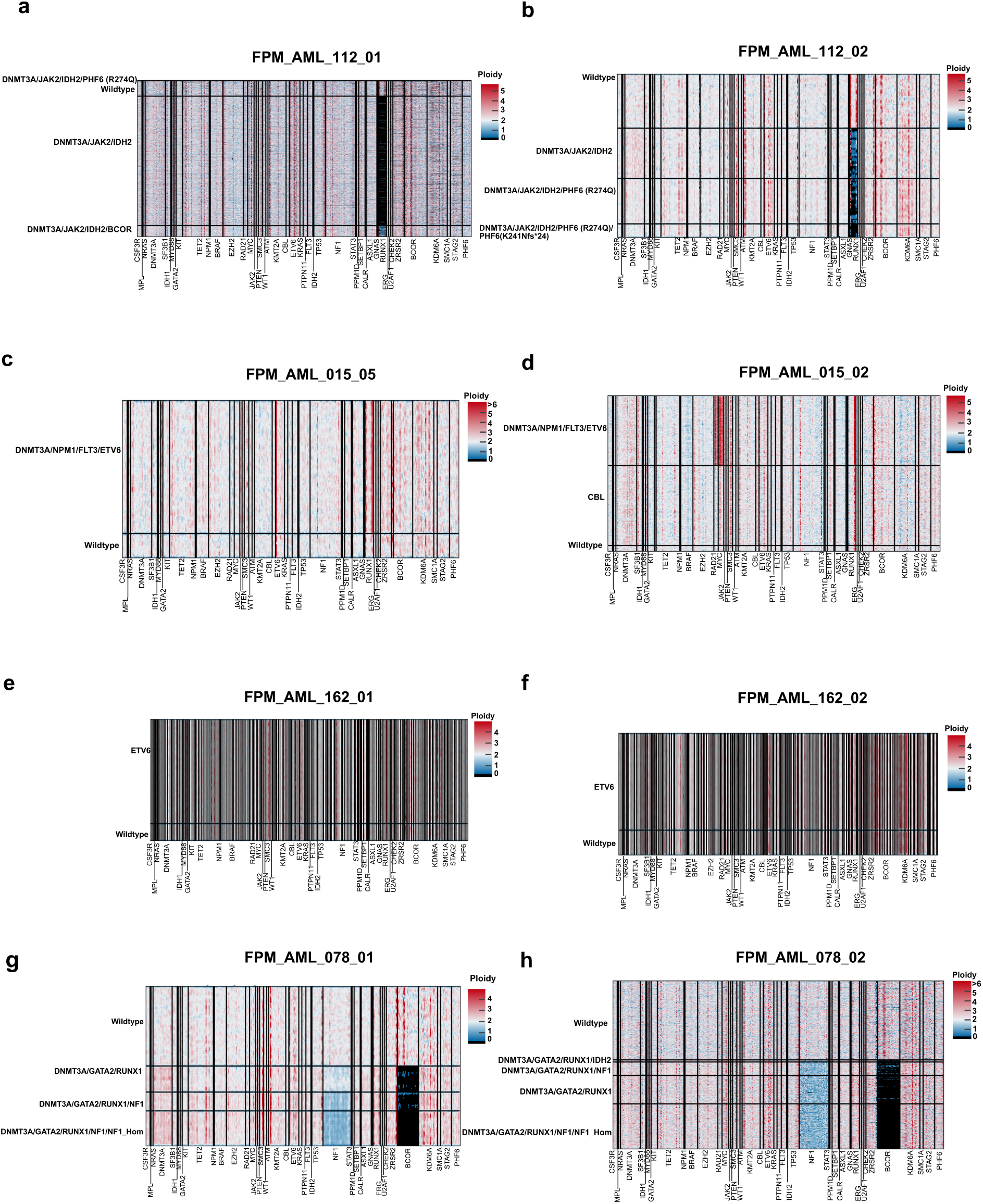
Gene-copy number profile for eight AML samples of genes detected by scDNA-seq. Each dot on the x-axis represents a cell and the yaxis shows amplicons. The values have been smoothed by a moving average.

**Supplementary Figure 2.**
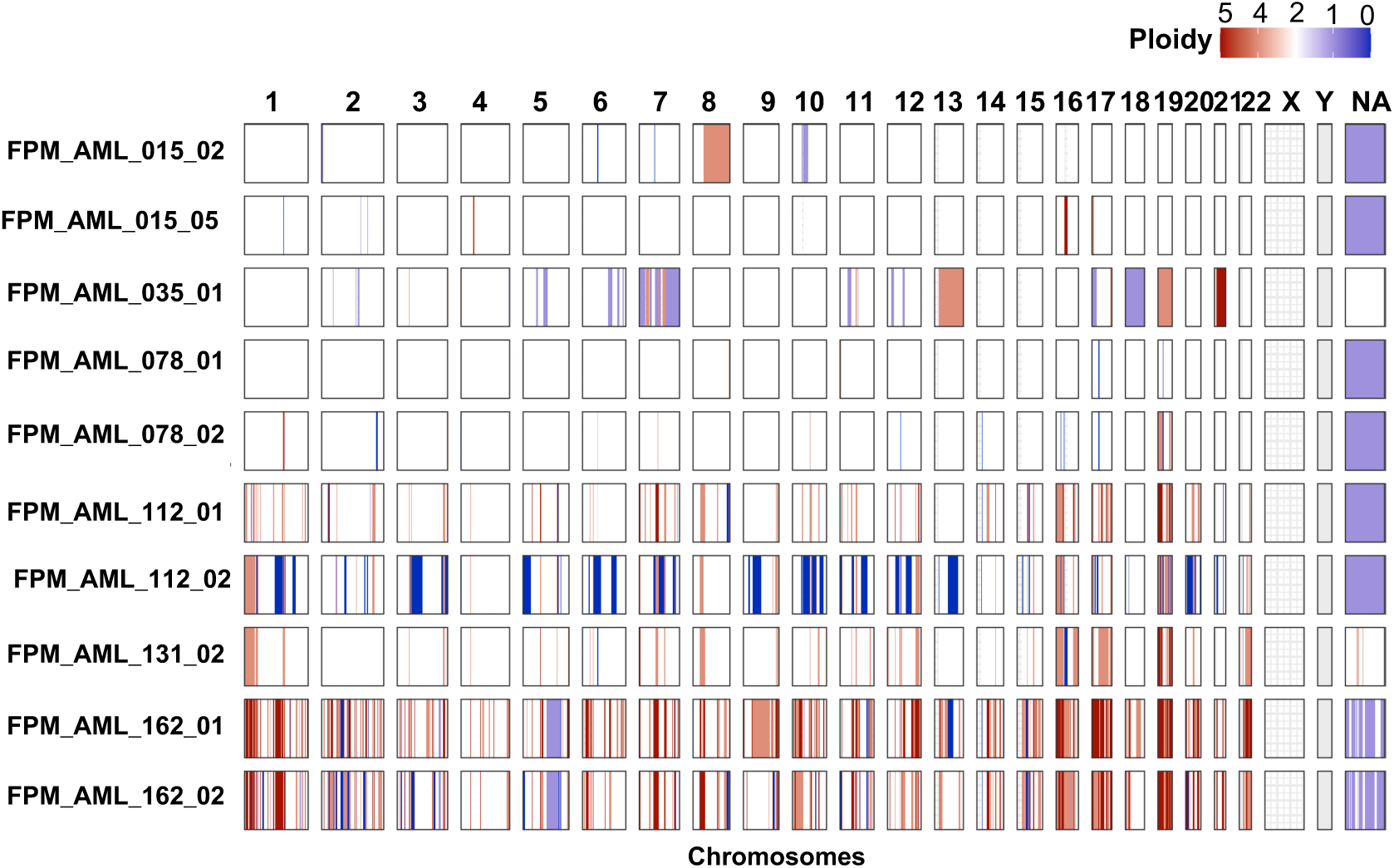
Genome-wide copy numb shows the chromosomes and y-axis shows the sampl produces chromosome 23 output instead of X and Y, shown as NA.

**Supplementary Figure 3.**
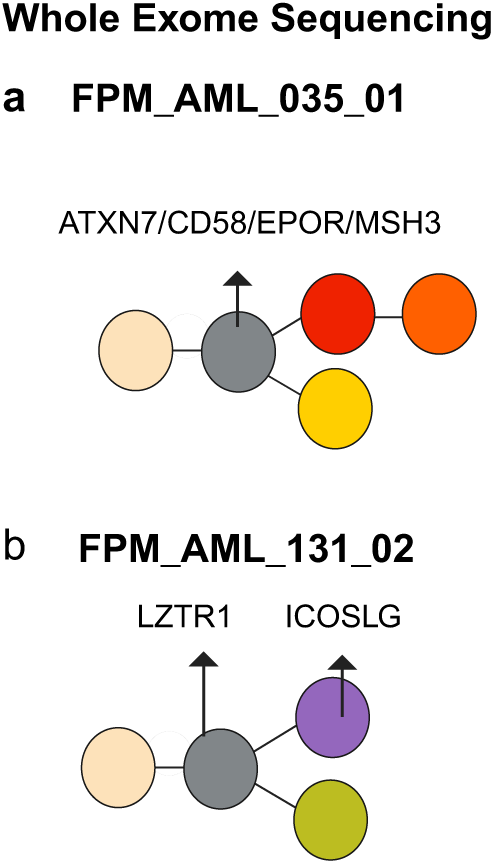
Phylogeny for sample FPM_AML_035_01 (a) and FPM_AML_131_02 (b) depicted by WES.

