## Supplementary tables for "Whole Exome Sequencing and Single-Cell DNA Sequencing for Assessment of Clonal Heterogeneity and Evolution in Acute Myeloid Leukemia"

Supplementary Table 1. Quality statistics for WES (top) and scDNA-seq (bottom)

| Whole Exome Sequencing | | | | |
|---|---|---|---|---|
| Sample | Total reads | Mapped reads | Reads mapped to target | Number of somatic variants |
| FPM_AML_112_01 | 118242623 | 98.49% | 61.04% | 36 |
| FPM_AML_112_02 | 133731007 | 98.86% | 80.83% | 94 |
| FPM_AML_131_02 | 134542926 | 99.15% | 81.9% | 100 |
| FPM_AML_162_01 | 146009624 | 98.5% | 77.09% | 61 |
| FPM_AML_162_02 | 107791282 | 99.08% | 77.9% | 58 |
| FPM_AML_015_05 | 274941606 | 96.6% | 40.06% | 24 |
| FPM_AML_015_02 | 102121840 | 96.11% | 39.05% | 180 |
| FPM_AML_078_01 | 89234052 | 99.03% | 79.03% | 37 |
| FPM_AML_078_02 | 93052142 | 98.47% | 71.97% | 61 |
| FPM_AML_035_01 | 64506901 | 99.08% | 70.26% | 41 |
| Mean | 126417400 | 98.34% | 67.91% | 69,2 |
| Total | 1264174003 | - | - | 692 |
| median | 113016952 | 98.68% | 74.53% | 59,5 |

| Single cell DNA sequencing | | | | | | | | |
|---|---|---|---|---|---|---|---|---|
| Sample | Number of cells | Mean reads per cell | Mean reads per cell per amplicon | Reads mapped to target | ADO rate | Total number of all variants | Filtered Number of cells | Filtered Number of Variants |
| FPM_AML_112_01 | 7830 | 20693 | 66 | 88.97% | 10.60% | 107 | 2233 | 4 |
| FPM_AML_112_02 | 6602 | 32570 | 104 | 89.38% | 10.50% | 112 | 1186 | 5 |
| FPM_AML_131_02 | 9726 | 24779 | 79 | 89.22 % | 13.80% | 86 | 0 | 0 |
| FPM_AML_162_01 | 5190 | 40819 | 130 | 90.10 % | 10.20% | 107 | 4211 | 1 |
| FPM_AML_162_02 | 5154 | 47183 | 151 | 81.55 % | 13.20% | 115 | 4532 | 1 |
| FPM_AML_015_05 | 7048 | 23072 | 73 | 87.65 % | 13.10% | 114 | 633 | 5 |
| FPM_AML_015_02 | 7833 | 20754 | 66 | 89.93 % | 8.05% | 122 | 1817 | 6 |
| FPM_AML_078_01 | 6710 | 22570 | 72 | 87.80 % | 11.00% | 87 | 1158 | 4 |
| FPM_AML_078_02 | 8546 | 25424 | 81 | 87.19 % | 16.50% | 95 | 1284 | 5 |
| FPM_AML_035_01 | 8851 | 17250 | 55 | 90.66 % | 19.20% | 79 | 0 | 0 |
| Mean | 7349 | 28651,5556 | 91,3333333 | 88.245% | 12.615% | 102,4 | 1705,4 | 3.1 |
| Total | 73490 | 257864 | 822 | - | - | 1024 | 17054 | 31 |
| median | 7439 | 24779 | | 88.97 | 12.05% | 107 | 1235 | 4 |

Supplementary Table 2: Genes included in the Tapestri myeloid panel

| Chromosome | Genes |
|---|---|
| chr1 | CSF3R, MPL,NRAS |
| chr2 | DNMT3A, SF3B1, IDH1 |
| chr3 | MYD88, GATA2 |
| chr4 | KIT, TET2 |
| chr5 | NPM1 |
| chr6 | |
| chr7 | BRAF, EZH2 |
| chr8 | RAD21, MYC |
| chr9 | JAK2 |
| chr10 | PTEN, SMC3 |
| chr11 | WT1, ATM, KMT2A, CBL |
| chr12 | ETV6, KRAS, PTPN11 |
| chr13 | FLT3 |
| chr14 | |
| chr15 | IDH2 |
| chr16 | |
| chr17 | NF1, STAT3, PPM1D, TP53 |
| chr18 | SETBP1 |
| chr19 | CALR |
| chr20 | ASXL1, GNAS |
| chr21 | SETBP1 |
| chr22 | CHEK2 |
| chrX | ZRSR2, BCOR, KDM6A, SMC1A, STAG2,PHF6 |

Supplementary Table 3: Clones identified in sample FPM_AML_112 via scDNA-seq using a WES-derived, whitelisted ASXL1 variant position.

| Variant | ASXL1:chr20:31022441:A/AG | BCOR:chrX:39916415:C/A | DNMT3A:chr2:25467449:C/A | IDH2:chr15:90631934:C/T | JAK2:chr9:5073770:G/T | PHF6:chrX:133547989:A/AC | PHF6:chrX:133549137:G/A | Total | FPM_AML_112_01.cells | FPM_AML_112_02.cells |
|---|---|---|---|---|---|---|---|---|---|---|
| C1 | WT (1%) | WT (0%) | Het (53%) | Het (51%) | Het (53%) | WT (0%) | WT (1%) | 466 (3.23%) | 349 (4.46%) | 117 (1.77%) |
| C2 | WT (3%) | WT (0%) | Het (52%) | Het (52%) | Het (54%) | WT (1%) | Hom (99%) | 169 (1.17%) | 0 (0.00%) | 169 (2.56%) |
| WT | WT (3%) | WT (0%) | WT (0%) | WT (0%) | WT (0%) | WT (0%) | WT (1%) | 140 (0.97%) | 51 (0.65%) | 89 (1.35%) |
| C4 | WT (3%) | WT (0%) | Het (52%) | Het (51%) | Het (51%) | WT (0%) | Het (58%) | 95 (0.66%) | 0 (0.00%) | 95 (1.44%) |
| C5 | WT (1%) | Het (54%) | Het (50%) | Het (52%) | Het (48%) | WT (0%) | WT (1%) | 45 (0.31%) | 45 (0.57%) | 0 (0.00%) |
| C6 | WT (2%) | WT (0%) | Het (55%) | Het (55%) | Hom (100%) | WT (0%) | WT (1%) | 39 (0.27%) | 15 (0.19%) | 24 (0.36%) |
| C7 | WT (4%) | WT (0%) | Het (54%) | Het (49%) | Het (49%) | Het (48%) | Het (54%) | 33 (0.23%) | 0 (0.00%) | 33 (0.50%) |
| C8 | WT (2%) | WT (0%) | Het (46%) | Het (51%) | WT (1%) | WT (0%) | WT (1%) | 32 (0.22%) | 23 (0.29%) | 9 (0.14%) |
| C9 | WT (2%) | WT (0%) | Het (47%) | WT (0%) | Het (53%) | WT (0%) | WT (1%) | 28 (0.19%) | 18 (0.23%) | 10 (0.15%) |
| C10 | WT (1%) | WT (0%) | Hom (99%) | Het (59%) | Het (53%) | WT (0%) | WT (1%) | 26 (0.18%) | 18 (0.23%) | 8 (0.12%) |
| C11 | WT (1%) | Hom (100%) | Het (49%) | Het (52%) | Het (50%) | WT (0%) | WT (1%) | 25 (0.17%) | 25 (0.32%) | 0 (0.00%) |
| C12 | WT (3%) | WT (0%) | Het (48%) | Het (49%) | WT (1%) | WT (0%) | Hom (99%) | 20 (0.14%) | 0 (0.00%) | 20 (0.30%) |
| C13 | WT (2%) | WT (0%) | WT (1%) | Het (48%) | Het (56%) | WT (0%) | WT (1%) | 19 (0.13%) | 16 (0.20%) | 3 (0.05%) |
| C14 | WT (3%) | WT (0%) | Het (54%) | Het (58%) | Het (49%) | Hom (99%) | WT (0%) | 18 (0.12%) | 0 (0.00%) | 18 (0.27%) |
| C15 | WT (3%) | WT (0%) | Het (47%) | WT (0%) | WT (0%) | WT (0%) | WT (1%) | 14 (0.10%) | 4 (0.05%) | 10 (0.15%) |
| C16 | WT (2%) | WT (1%) | Het (59%) | Het (60%) | Hom (98%) | WT (0%) | Hom (99%) | 14 (0.10%) | 0 (0.00%) | 14 (0.21%) |
| C17 | Het (30%) | WT (0%) | Het (59%) | Het (60%) | Het (51%) | WT (0%) | WT (1%) | 13 (0.09%) | 6 (0.08%) | 7 (0.11%) |
| C18 | WT (3%) | WT (0%) | Het (56%) | Hom (99%) | Het (52%) | WT (0%) | WT (1%) | 12 (0.08%) | 8 (0.10%) | 4 (0.06%) |
| C19 | WT (4%) | WT (1%) | Het (56%) | WT (1%) | Het (55%) | WT (0%) | Hom (99%) | 10 (0.07%) | 0 (0.00%) | 10 (0.15%) |
| C20 | WT (3%) | WT (0%) | WT (0%) | Het (48%) | Het (52%) | WT (1%) | Hom (99%) | 9 (0.06%) | 0 (0.00%) | 9 (0.14%) |
| C21 | WT (2%) | WT (0%) | Het (48%) | Hom (99%) | Het (57%) | WT (0%) | Hom (100%) | 9 (0.06%) | 0 (0.00%) | 9 (0.14%) |
| C22 | WT (3%) | WT (0%) | Hom (98%) | Het (45%) | Het (63%) | WT (0%) | Hom (100%) | 7 (0.05%) | 0 (0.00%) | 7 (0.11%) |
| C23 | WT (2%) | WT (0%) | Het (54%) | Het (56%) | Het (55%) | Het (53%) | WT (2%) | 7 (0.05%) | 0 (0.00%) | 7 (0.11%) |
| C24 | WT (1%) | WT (0%) | Het (52%) | WT (0%) | Hom (98%) | WT (0%) | WT (1%) | 6 (0.04%) | 2 (0.03%) | 4 (0.06%) |
| C25 | WT (4%) | WT (0%) | WT (0%) | Het (48%) | Hom (100%) | WT (1%) | WT (1%) | 5 (0.03%) | 1 (0.01%) | 4 (0.06%) |
| C26 | WT (4%) | WT (0%) | Het (42%) | WT (1%) | Het (70%) | Hom (100%) | WT (0%) | 4 (0.03%) | 0 (0.00%) | 4 (0.06%) |
| C27 | WT (1%) | WT (0%) | WT (2%) | Het (66%) | Het (46%) | Hom (100%) | WT (1%) | 4 (0.03%) | 0 (0.00%) | 4 (0.06%) |
| C28 | WT (2%) | WT (0%) | WT (1%) | WT (0%) | Het (66%) | WT (0%) | WT (0%) | 4 (0.03%) | 1 (0.01%) | 3 (0.05%) |
| C29 | WT (0%) | WT (0%) | Het (45%) | Het (45%) | WT (3%) | Hom (99%) | WT (1%) | 3 (0.02%) | 0 (0.00%) | 3 (0.05%) |
| C30 | WT (5%) | WT (0%) | WT (2%) | Het (33%) | Het (37%) | WT (1%) | Het (49%) | 3 (0.02%) | 0 (0.00%) | 3 (0.05%) |
| C31 | WT (3%) | WT (0%) | WT (0%) | Hom (99%) | Het (69%) | WT (1%) | WT (1%) | 3 (0.02%) | 1 (0.01%) | 2 (0.03%) |
| C32 | WT (2%) | WT (0%) | Het (42%) | WT (0%) | WT (0%) | WT (0%) | Hom (99%) | 3 (0.02%) | 0 (0.00%) | 3 (0.05%) |
| C33 | Het (27%) | WT (0%) | Het (31%) | Het (47%) | Het (39%) | WT (1%) | Hom (99%) | 3 (0.02%) | 0 (0.00%) | 3 (0.05%) |
| C34 | Het (37%) | WT (0%) | Het (53%) | Het (47%) | Het (61%) | WT (1%) | Het (54%) | 3 (0.02%) | 0 (0.00%) | 3 (0.05%) |
| C35 | WT (2%) | WT (0%) | Het (39%) | Hom (100%) | Hom (98%) | WT (0%) | WT (1%) | 2 (0.01%) | 1 (0.01%) | 1 (0.02%) |
| C36 | WT (3%) | WT (1%) | WT (0%) | WT (1%) | Hom (100%) | WT (0%) | WT (1%) | 2 (0.01%) | 0 (0.00%) | 2 (0.03%) |
| C37 | WT (6%) | WT (0%) | WT (3%) | WT (0%) | Het (60%) | WT (0%) | Hom (99%) | 2 (0.01%) | 0 (0.00%) | 2 (0.03%) |
| C38 | WT (5%) | WT (0%) | Hom (97%) | Het (42%) | Het (69%) | WT (0%) | Het (43%) | 2 (0.01%) | 0 (0.00%) | 2 (0.03%) |
| C39 | WT (2%) | WT (0%) | WT (0%) | Het (52%) | WT (2%) | WT (1%) | WT (1%) | 2 (0.01%) | 0 (0.00%) | 2 (0.03%) |
| C40 | WT (3%) | WT (2%) | Hom (100%) | WT (0%) | WT (0%) | WT (0%) | WT (0%) | 2 (0.01%) | 0 (0.00%) | 2 (0.03%) |
| C41 | WT (0%) | WT (0%) | Het (67%) | WT (1%) | Hom (96%) | WT (1%) | Hom (100%) | 2 (0.01%) | 0 (0.00%) | 2 (0.03%) |
| C42 | WT (4%) | WT (0%) | Hom (99%) | Hom (100%) | Het (36%) | WT (1%) | WT (0%) | 2 (0.01%) | 1 (0.01%) | 1 (0.02%) |
| C43 | WT (1%) | WT (0%) | Hom (100%) | Hom (99%) | Het (62%) | WT (0%) | Hom (99%) | 2 (0.01%) | 0 (0.00%) | 2 (0.03%) |
| C44 | WT (1%) | WT (0%) | Het (62%) | Het (51%) | WT (1%) | WT (0%) | Het (49%) | 2 (0.01%) | 0 (0.00%) | 2 (0.03%) |
| C45 | WT (1%) | WT (0%) | Hom (98%) | Het (57%) | Hom (100%) | WT (1%) | WT (0%) | 2 (0.01%) | 0 (0.00%) | 2 (0.03%) |
| C46 | WT (0%) | Hom (100%) | Het (24%) | Het (58%) | Hom (98%) | WT (0%) | WT (2%) | 2 (0.01%) | 2 (0.03%) | 0 (0.00%) |
| C47 | Het (52%) | WT (1%) | WT (1%) | Het (52%) | Het (42%) | WT (0%) | WT (2%) | 2 (0.01%) | 2 (0.03%) | 0 (0.00%) |
| C48 | Het (31%) | WT (0%) | Het (46%) | Het (54%) | Hom (97%) | WT (0%) | WT (0%) | 2 (0.01%) | 0 (0.00%) | 2 (0.03%) |
| C49 | WT (6%) | WT (0%) | WT (1%) | Het (55%) | Hom (100%) | WT (0%) | Hom (100%) | 2 (0.01%) | 0 (0.00%) | 2 (0.03%) |
| C50 | WT (0%) | Hom (100%) | WT (0%) | Het (76%) | Het (38%) | WT (0%) | WT (0%) | 1 (0.01%) | 1 (0.01%) | 0 (0.00%) |
| C51 | WT (0%) | Hom (98%) | WT (0%) | Het (52%) | Hom (100%) | WT (0%) | WT (0%) | 1 (0.01%) | 1 (0.01%) | 0 (0.00%) |
| C52 | WT (0%) | Hom (100%) | Het (63%) | WT (0%) | Het (30%) | WT (0%) | WT (0%) | 1 (0.01%) | 1 (0.01%) | 0 (0.00%) |
| C53 | WT (0%) | Hom (100%) | Het (23%) | Het (77%) | WT (0%) | WT (0%) | WT (0%) | 1 (0.01%) | 1 (0.01%) | 0 (0.00%) |
| C54 | WT (6%) | Hom (96%) | Het (57%) | Hom (100%) | Hom (100%) | WT (0%) | WT (0%) | 1 (0.01%) | 1 (0.01%) | 0 (0.00%) |
| C55 | WT (0%) | Hom (100%) | Hom (100%) | WT (2%) | Het (29%) | WT (0%) | WT (0%) | 1 (0.01%) | 1 (0.01%) | 0 (0.00%) |
| C56 | WT (0%) | Het (54%) | WT (6%) | Het (85%) | WT (0%) | WT (0%) | WT (2%) | 1 (0.01%) | 1 (0.01%) | 0 (0.00%) |
| C57 | WT (0%) | Het (36%) | WT (5%) | Het (58%) | Het (38%) | WT (0%) | WT (0%) | 1 (0.01%) | 1 (0.01%) | 0 (0.00%) |
| C58 | WT (2%) | Het (62%) | Het (60%) | Het (50%) | WT (0%) | WT (0%) | WT (2%) | 1 (0.01%) | 1 (0.01%) | 0 (0.00%) |
| C59 | WT (4%) | Het (42%) | Het (52%) | Het (52%) | Het (55%) | WT (0%) | Het (33%) | 1 (0.01%) | 1 (0.01%) | 0 (0.00%) |
| C60 | WT (4%) | Hom (100%) | Hom (98%) | Het (63%) | Het (36%) | WT (0%) | WT (2%) | 1 (0.01%) | 1 (0.01%) | 0 (0.00%) |
| C61 | WT (0%) | WT (0%) | Het (48%) | Het (59%) | Hom (96%) | WT (0%) | Het (38%) | 1 (0.01%) | 0 (0.00%) | 1 (0.02%) |
| C62 | WT (0%) | WT (0%) | Het (27%) | Het (34%) | WT (2%) | Het (50%) | WT (3%) | 1 (0.01%) | 0 (0.00%) | 1 (0.02%) |
| C63 | WT (0%) | WT (0%) | Het (28%) | Het (52%) | Het (32%) | Het (82%) | Hom (96%) | 1 (0.01%) | 0 (0.00%) | 1 (0.02%) |
| C64 | WT (3%) | WT (0%) | Het (45%) | Hom (100%) | WT (0%) | WT (0%) | WT (0%) | 1 (0.01%) | 1 (0.01%) | 0 (0.00%) |
| C65 | WT (2%) | WT (0%) | Het (20%) | Hom (94%) | Het (36%) | Hom (100%) | WT (0%) | 1 (0.01%) | 0 (0.00%) | 1 (0.02%) |
| C66 | WT (0%) | WT (0%) | WT (0%) | Hom (100%) | Hom (100%) | WT (0%) | WT (0%) | 1 (0.01%) | 1 (0.01%) | 0 (0.00%) |
| C67 | WT (2%) | WT (0%) | WT (0%) | Het (23%) | WT (0%) | WT (2%) | Het (90%) | 1 (0.01%) | 0 (0.00%) | 1 (0.02%) |
| C68 | WT (0%) | WT (0%) | WT (0%) | Het (47%) | WT (0%) | WT (0%) | Hom (100%) | 1 (0.01%) | 0 (0.00%) | 1 (0.02%) |
| C69 | WT (9%) | WT (0%) | WT (0%) | Hom (100%) | Hom (95%) | WT (0%) | Hom (100%) | 1 (0.01%) | 0 (0.00%) | 1 (0.02%) |
| C70 | WT (2%) | WT (0%) | Het (38%) | WT (3%) | WT (5%) | WT (0%) | Het (73%) | 1 (0.01%) | 0 (0.00%) | 1 (0.02%) |
| C71 | WT (0%) | WT (0%) | Het (82%) | Hom (100%) | Hom (100%) | Hom (100%) | WT (0%) | 1 (0.01%) | 0 (0.00%) | 1 (0.02%) |
| C72 | WT (5%) | WT (4%) | Hom (100%) | Het (71%) | Hom (100%) | WT (2%) | Hom (98%) | 1 (0.01%) | 0 (0.00%) | 1 (0.02%) |
| C73 | WT (5%) | WT (0%) | Hom (100%) | Het (62%) | Het (40%) | Hom (100%) | WT (0%) | 1 (0.01%) | 0 (0.00%) | 1 (0.02%) |
| C74 | WT (0%) | WT (0%) | Hom (100%) | Het (26%) | WT (0%) | WT (0%) | Hom (100%) | 1 (0.01%) | 0 (0.00%) | 1 (0.02%) |
| C75 | WT (4%) | WT (5%) | Hom (100%) | WT (2%) | Het (45%) | WT (0%) | Hom (100%) | 1 (0.01%) | 0 (0.00%) | 1 (0.02%) |
| C76 | WT (0%) | WT (2%) | Hom (94%) | WT (0%) | Het (42%) | WT (0%) | WT (2%) | 1 (0.01%) | 0 (0.00%) | 1 (0.02%) |
| C77 | Het (20%) | WT (4%) | Het (84%) | Hom (96%) | Het (31%) | WT (0%) | WT (5%) | 1 (0.01%) | 1 (0.01%) | 0 (0.00%) |
| C78 | Het (31%) | WT (0%) | Hom (100%) | Het (40%) | Het (69%) | WT (0%) | Hom (100%) | 1 (0.01%) | 0 (0.00%) | 1 (0.02%) |
| C79 | Het (27%) | Hom (100%) | Het (65%) | Het (26%) | WT (0%) | WT (0%) | WT (0%) | 1 (0.01%) | 1 (0.01%) | 0 (0.00%) |
| C80 | Het (21%) | Hom (100%) | Het (60%) | Het (71%) | Het (31%) | WT (0%) | WT (0%) | 1 (0.01%) | 1 (0.01%) | 0 (0.00%) |
| C81 | Het (22%) | WT (0%) | Het (35%) | Het (45%) | Hom (100%) | WT (0%) | Hom (100%) | 1 (0.01%) | 0 (0.00%) | 1 (0.02%) |
| C82 | Het (21%) | WT (0%) | WT (4%) | Het (65%) | Het (45%) | Het (50%) | Het (58%) | 1 (0.01%) | 0 (0.00%) | 1 (0.02%) |
| C83 | Het (31%) | WT (0%) | WT (2%) | Hom (100%) | WT (0%) | WT (2%) | WT (2%) | 1 (0.01%) | 0 (0.00%) | 1 (0.02%) |
| C84 | Het (20%) | WT (0%) | Het (21%) | WT (0%) | Het (72%) | WT (2%) | Het (24%) | 1 (0.01%) | 0 (0.00%) | 1 (0.02%) |
| C85 | Het (22%) | WT (0%) | Het (44%) | Het (73%) | Het (54%) | Het (50%) | WT (0%) | 1 (0.01%) | 0 (0.00%) | 1 (0.02%) |
| C86 | Het (21%) | WT (0%) | Het (54%) | Het (69%) | Het (41%) | Hom (100%) | WT (2%) | 1 (0.01%) | 0 (0.00%) | 1 (0.02%) |
| Small Subclor | - | - | - | - | - | - | - | 0 (0.00%) | 0 (0.00%) | 0 (0.00%) |
| Missing GT S | Missing in 81.77% of clones | Missing in 21.46% of clones | Missing in 11.07% of clones | Missing in 11.97% of clones | Missing in 21.50% of clones | Missing in 4.82% of clones | Missing in 4.95% of clones | 13074 (90.59 | 7225 (92.27%) | 5849 (88.59%) |

Supplementary Table 4. Details of clonal composition, including cellular prevalence and Shannon index. Additionally, for WES the number of mutations in each clone is included. If driver mutations were not present, only the number of mutations is shown.

| **WES** | | | | |
|---|---|---|---|---|
| Patient | Clones and Number_of_mutation | Sample | Cellular_Prevalance | Shannon_index |
| FPM_AML_112 | DNMT3A/IDH2/JAK2/ASXL1/PTPRS/ZNRF3 | FPM_AML_112_01 | 80 | 1.17 |
| | LRP1B n=13 | FPM_AML_112_01 | 49 | |
| | n=25 | FPM_AML_112_01 | 20 | |
| | BRAF n=12 | FPM_AML_112_01 | 0 | |
| | BCOR n=2 | FPM_AML_112_01 | 13 | |
| | PHF6 n=4 | FPM_AML_112_01 | 0 | |
| | n=31 | FPM_AML_112_01 | 0 | |
| | DNMT3A/IDH2/JAK2/ASXL1/PTPRS/ZNRF3 | FPM_AML_112_02 | 82 | |
| | LRP1B n=13 | FPM_AML_112_02 | 37 | |
| | n=25 | FPM_AML_112_02 | 19 | |
| | BRAF n=12 | FPM_AML_112_02 | 11 | |
| | BCOR n=2 | FPM_AML_112_02 | 0 | 1.41 |
| | PHF6 n=4 | FPM_AML_112_02 | 43 | |
| | n=31 | | | |
| FPM_AML_162 | ETV6/MAP2K1/PGBD5 n=21 | FPM_AML_162_01 | 82 | 1.62 |
| | AR n=6 | FPM_AML_162_01 | 54 | |
| | ICOSLG n=19 | FPM_AML_162_01 | 12 | |
| | ICOSLG/SDHA n=8 | FPM_AML_162_01 | 12 | |
| | n=1 | FPM_AML_162_01 | 16 | |
| | n=28 | FPM_AML_162_01 | 19 | |
| | n=4 | FPM_AML_162_01 | 6 | |
| | FOXP1 n=3 | FPM_AML_162_01 | 2 | |
| | n=2 | FPM_AML_162_01 | 0 | |
| | ETV6/MAP2K1/PGBD5 n=21 | FPM_AML_162_02 | 83 | 1.78 |
| | AR n=6 | FPM_AML_162_02 | 26 | |
| | ICOSLG n=19 | FPM_AML_162_02 | 12 | |
| | ICOSLG/SDHA n=8 | FPM_AML_162_02 | 7 | |
| | n=1 | FPM_AML_162_02 | 0 | |
| | n=28 | FPM_AML_162_02 | 25 | |
| | n=4 | FPM_AML_162_02 | 14 | |
| | FOXP1 n=3 | FPM_AML_162_02 | 27 | |
| | n=2 | FPM_AML_162_02 | 12 | |
| FPM_AML_015 | DNMT3A/ETV6/FLT3/NPM1 n=17 | FPM_AML_015_05 | 77 | 1.22 |
| | EP400 n=8 | FPM_AML_015_05 | 47 | |
| | FLT3 n=3 | FPM_AML_015_05 | 13 | |
| | ACKR3/CBL/ERCC4/FOLH1/MTOR/PMS1/ | FPM_AML_015_05 | 0 | |
| | ARID3A/FANCD2/FOLH1/USP8 n=24 | FPM_AML_015_05 | 0 | |
| | n=6 | FPM_AML_015_05 | 15 | |
| | USP8 n=14 | FPM_AML_015_05 | 3 | |
| | DNMT3A/ETV6/FLT3/NPM1 n=17 | FPM_AML_015_02 | 72 | 1.49 |
| | EP400 n=8 | FPM_AML_015_02 | 0 | |
| | FLT3 n=3 | FPM_AML_015_02 | 0 | |
| | ACKR3/CBL/ERCC4/FOLH1/MTOR/PMS1/ | FPM_AML_015_02 | 42 | |
| | ARID3A/FANCD2/FOLH1/USP8 n=24 | FPM_AML_015_02 | 21 | |
| | n=6 | FPM_AML_015_02 | 26 | |
| | USP8 n=14 | FPM_AML_015_02 | 22 | |
| FPM_AML_078 | DNMT3A/GATA2 n=20 | FPM_AML_078_01 | 72 | 1.62 |
| | RUNX1 n=1 | FPM_AML_078_01 | 49 | |
| | n=2 | FPM_AML_078_01 | 29 | |
| | n=4 | FPM_AML_078_01 | 15 | |
| | NF1 n=15 | FPM_AML_078_01 | 13 | |
| | n=14 | FPM_AML_078_01 | 2 | |
| | n=15 | FPM_AML_078_01 | 15 | |
| | DNMT3A/GATA2 n=20 | FPM_AML_078_02 | 72 | 1.45 |
| | RUNX1 n=1 | FPM_AML_078_02 | 40 | |
| | n=2 | FPM_AML_078_02 | 0 | |
| | n=4 | FPM_AML_078_02 | 0 | |
| | NF1 n=15 | FPM_AML_078_02 | 16 | |
| | n=14 | FPM_AML_078_02 | 18 | |
| | n=15 | FPM_AML_078_02 | 26 | |
| FPM_AML_131 | LZTR1 n=15 | FPM_AML_131_02 | 85 | 1.20 |
| | n=18 | FPM_AML_131_02 | 53 | |
| | ICOSLG n=55 | FPM_AML_131_02 | 15 | |
| | n=11 | FPM_AML_131_02 | 28 | |
| FPM_AML_035 | ATXN7/CD58/EPOR/MSH3 n=32 | FPM_AML_035_01 | 74 | 0.94 |
| | n=4 | FPM_AML_035_01 | 40 | |
| | n=3 | FPM_AML_035_01 | 17 | |

| **ScDNA-seq** | | | | |
|---|---|---|---|---|
| Patient | Clone | Sample | Cellular_Prev | Shannon_index |
| FPM_AML_112 | DNMT3A/JAK2/IDH2 | FPM_AML_112_01 | 89 | 0.257 |
| | DNMT3A/JAK2/IDH2/PHF6(R274Q) | FPM_AML_112_01 | 0.1 | |
| | DNMT3A/JAK2/IDH2/PHF6(R274Q)/ | FPM_AML_112_01 | 0 | |
| | DNMT3A/JAK2/IDH2/BCOR | FPM_AML_112_01 | 6.5 | |
| | DNMT3A/JAK2/IDH2 | FPM_AML_112_02 | 67.2 | 0.86 |
| | DNMT3A/JAK2/IDH2/PHF6(R274Q) | FPM_AML_112_02 | 36 | |
| | DNMT3A/JAK2/IDH2/PHF6(R274Q)/ | FPM_AML_112_02 | 8.1 | |
| | DNMT3A/JAK2/IDH2/BCOR | FPM_AML_112_02 | 0 | |
| FPM_AML_162 | ETV6 | FPM_AML_162_01 | 84.84 | 0 |
| | ETV6 | FPM_AML_162_02 | 79.90 | 0 |
| FPM_AML_015 | DNMT3A/NPM1/FLT3/ETV6 | FPM_AML_015_05 | 84.49 | 0 |
| | CBL | FPM_AML_015_05 | 0 | |
| | DNMT3A/NPM1/FLT3/ETV6 | FPM_AML_015_02 | 43.52 | 0.689 |
| | CBL | FPM_AML_015_02 | 52.21 | |
| FPM_AML_078 | DNMT3A/GATA2/RUNX1 | FPM_AML_078_01 | 50.4 | '1.04 |
| | DNMT3A/GATA2/RUNX1/NF1 | FPM_AML_078_01 | 33.7 | |
| | DNMT3A/GATA2/RUNX1/NF1_Hom | FPM_AML_078_01 | 21.88 | |
| | DNMT3A/GATA2/RUNX1/IDH2 | FPM_AML_078_01 | 0 | |
| | DNMT3A/GATA2/RUNX1 | FPM_AML_078_02 | 53.73 | 1.1 |
| | DNMT3A/GATA2/RUNX1/NF1 | FPM_AML_078_02 | 34.8 | |
| | DNMT3A/GATA2/RUNX1/NF1_Hom | FPM_AML_078_02 | 26.79 | |
| | DNMT3A/GATA2/RUNX1/IDH2 | FPM_AML_078_02 | 1.26 | |
